# Alphavirus nsP3 ADP-ribosylhydrolase Activity Disrupts Stress Granule Formation

**DOI:** 10.1101/629881

**Authors:** Aravinth Kumar Jayabalan, Diane E. Griffin, Anthony K. L. Leung

## Abstract

Formation of stress granules (SGs), cytoplasmic condensates of stalled translation initiation complexes, is regulated by post-translational protein modification. Alphaviruses interfere with SG formation in response to inhibition of host protein synthesis through the activities of nonstructural protein 3 (nsP3). nsP3 has a conserved N-terminal macrodomain that binds and can remove ADP-ribose from ADP-ribosylated proteins and a C-terminal hypervariable domain that binds essential SG component G3BP1. We showed that the hydrolase activity of chikungunya virus nsP3 macrodomain removed ADP-ribosylation of G3BP1 and suppressed SG formation. ADP-ribosylhydrolase-deficient nsP3 mutants allowed stress-induced cytoplasmic condensation of translation initiation factors. nsP3 also disassembled SG-like aggregates enriched with translation initiation factors that are induced by the expression of FUS mutant R495X linked to amyotrophic lateral sclerosis. Therefore, our data indicate that regulation of ADP-ribosylation controls the localization of translation initiation factors during virus infection and other pathological conditions.

## INTRODUCTION

Non-membranous structures are prevalent in cells and critical for cellular functions, including RNA metabolism, embryonic cell fate specification, and neuronal activities [1–3]. Although the components of these non-membranous structures are often dynamically exchanged with the surrounding milieu, the compositions of these cellular structures remain distinct [4]. It is, however, unclear how individual components are selectively retained in these non-membranous structures. Stress granules (SGs), one of the best characterized dynamic non-membranous structures, are RNA–protein assemblies formed in response to a variety of environmental cues [3]. In most cases, these environmental cues activate stress-responsive protein kinases that phosphorylate the eukaryotic initiation factor 2 alpha (eIF2α), resulting in the stalling of translation initiation [3]. The sudden influx of untranslated mRNAs is proposed to seed the formation of SGs, where the polynucleotide promotes local concentration of proteins through non-covalent binding [5]. These RNA-binding proteins are highly enriched with low-complexity regions, and emergent data indicate that the nonspecific, weak interactions between these regions are responsible for the condensation of proteins to form higher-order structures, such as microscopically visible SGs [6–8]. Depending on the type of stress, the composition of SGs could vary [9], but certain common components, such as Ras GTP-activating protein-binding proteins G3BP1/2, are essential for SG formation [10,11]. Dysregulation of SG assembly/disassembly and mutations in the low-complexity region of specific SG proteins are implicated in the pathogenesis of diseases such as viral infection, cancer and neurodegeneration [1,12–14]. Therefore, understanding the regulatory mechanisms of SG assembly is critical for designing novel therapeutics.

SG assembly is regulated positively and negatively by post-translational modifications of proteins, including those that conjugate simple chemical groups (such as phosphorylation, O-GlcNAc glycosylation, methylation, acetylation), attach polypeptides (e.g., neddylation), and add nucleotides as in the case of ADP-ribosylation [15–21]. ADP-ribosylation refers to the addition of one or more ADP-ribose units onto proteins [22–24]. In humans, ADP-ribosylation is accomplished primarily by a family of 17 ADP-ribosyltransferases, a subset of which is commonly known as poly(ADP-ribose) polymerases (PARPs) [22–24]. Polymers of ADP-ribose [i.e., poly(ADP-ribose or PAR], five PARPs and two isoforms of the degradative enzyme PAR glycohydrolase (PARG) have been identified in SGs, where selective SG proteins are ADP-ribosylated [19,25,26]. Overexpression of these PARPs and PARG isoforms induces and suppresses SG formation, respectively, while PARG knockdown delays SG disassembly [19,25]. PAR, like RNA, has been proposed to seed formation of non-membranous structures by facilitating the high concentration of low complexity region-containing proteins locally through noncovalent binding to the repetitive monomeric units of the polymer [19,25–28]. The non-covalent PAR-protein interaction also facilitates the targeting of specific proteins to SGs [25,26,29], such as TDP-43—a key protein involved in several neurodegenerative diseases, including amyotrophic lateral sclerosis (ALS). Such a targeting mechanism based on PAR-binding to protein is proposed to regulate TDP-43 localization and prevent the formation of pathological aggregates in ALS patients [25,26]. Certain PARP inhibitors prevent the neurotoxicity in neuronal cells [26,30], suggesting that pharmacological modulation of PAR-mediated protein targeting could have potential therapeutic benefit.

SG assembly and disassembly are tightly regulated during infection by many viruses, often reflecting cellular translation status [31–34]. In the early phase of infection, the presence of double-stranded viral RNAs activates protein kinase R (PKR), resulting in eIF2α phosphorylation, mRNA translation stalling and assembly of SGs enriched with translation factors [32,34]. Yet, many viruses disassemble SGs or suppress SG formation [32,34], but the mechanisms and the physiological roles remain largely unclear. For example, the mosquito-borne alphaviruses, which cause a range of diseases from rashes and arthritis to encephalitis, induce SG formation transiently in the early stage of infection, followed by disassembly [32,34–36]. Previous studies have identified the alphaviral nonstructural protein 3 (nsP3), a key factor for virus replication and virulence [37–39], as able to suppress SG assembly [36,40–42]. nsP3 is a tripartite protein composed of a highly conserved <u>m</u>acro<u>d</u>omain (MD) in the N-terminus, a central zinc-binding domain, and a C-terminal <u>h</u>yper<u>v</u>ariable <u>d</u>omain (HVD), which is of low-complexity [39]. Recent studies indicate that short peptide motifs within the HVD direct nsP3 binding to host SG proteins [39,40,43]. Specifically, two repeat peptide motifs within the HVD of the alphavirus chikungunya virus (CHIKV) bind to the essential SG components G3BP1 and G3BP2 [40,44]. Given that the expression of alphaviral nsP3 alone suppresses SG formation and nsP3 expression increases over the course of viral infection, it was proposed that nsP3 sequesters G3BP1/2 resulting in SG disassembly during the late phase of infection [35,36,41].

Here, we report that the expression of the G3BP-binding HVD alone did not suppress SG formation; instead, the expression of the N-terminal MD alone is sufficient to suppress SG formation. Recently, we and others have reported that the viral MD possesses enzymatic activity for removal of single ADP-ribose groups, and possibly PAR, from ADP-ribosylated proteins [38,45–47]. Given that the structural integrity of SGs is dependent on ADP-ribosylation [19], we hypothesized, and identified, that the MD ADP-ribosylhydrolase activity is required for suppressing the formation of SGs induced by stress, with G3BP1 as one of the target substrates. This enzymatic activity is required for alteration of SG composition by releasing translation factors from a condensated state in nsP3-expressing cells. Intriguingly, nsP3 expression also suppresses the formation of and disassembles SG-like aggregates induced by a pathological mutant of the RNA-binding protein FUS (R495X) from an aggressive form of ALS [48,49]. These data altogether suggest that nsP3 ADP-ribosylhydrolase activity is able to disrupt SG formation in both physiological and pathological conditions.

## RESULTS

### The hypervariable domain recruits nsP3 to SGs and the macrodomain suppresses SG formation

Overexpression of alphaviral nsP3 alone suppresses the formation of SGs even in the presence of the classic SG inducer arsenite [36,40–42]. The current paradigm is that the essential SG component G3BP1 is sequestered by nsP3 in another class of cytoplasmic structure that does not contain the canonical SG marker translation initiation factor eIF3b (Fig. 1a). Given that G3BP1 and its paralog G3BP2 bind to the C-terminal HVD of nsP3 (nsP3^HVD^) [40,41,44,50–52], their localization in these nsP3-positive foci is expected. However, it is unclear whether the mechanism for sequestration of G3BP by the HVD is sufficient to cause the disappearance of SGs because the domain binds multiple cellular proteins [39,50,53] and cellular G3BP1/2 are highly abundant (Table S1).

**Figure 1.**
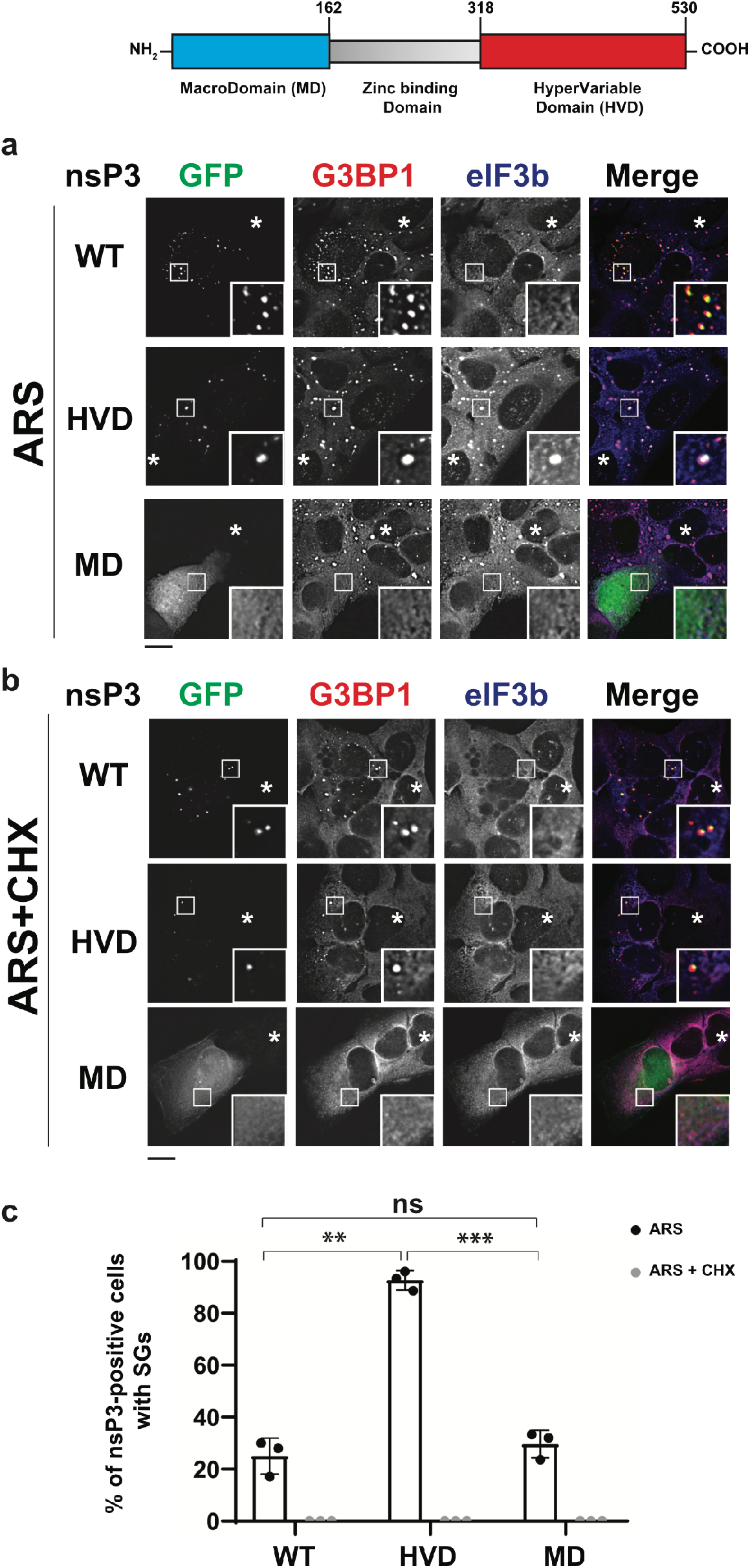
nsP3 macrodomain suppresses SG formation. Schematic representation of the domain architecture of nsP3. U20S cells transfected with GFP-tagged WT nsP3, nsP3^HvD^ or nsP3^MD^ were treated with (a) 0.2 mM arsenite (ARS) or (b) 0.2 mM arsenite with 100 μg/ml cycloheximide (ARS+CHX) for 30 min. Coverslips were processed and immunostained for SG markers elF3b (blue) and G3BP1 (red). Asterisks indicate untransfected cells. (c) Bar graph shows the percentage of nsP3-positive cells with SGs. **, p<0.01, ***, p<0.001, two tailed, unpaired Student’s t test. Scale bar, 10 μm.

To determine whether the nsP3^HVD^ alone is sufficient to suppress arsenite-induced SG formation, we overexpressed either GFP-tagged full-length nsP3 or the HVD fragment (318-530) and examined the co-localization of nsP3 with SG components G3BP1 and eIF3b in U2OS (Fig. 1a, S1a) and HeLa cells (Fig. S1b). In untransfected cells, eIF3b and G3BP1 co-localized as foci (i.e., SGs) after arsenite treatment (Fig. 1a, asterisk). In cells transfected with full-length nsP3, nsP3 co-localized with G3BP1, but not with eIF3b in the majority of cells examined (73%; Fig. 1a, c), as previously reported [36,42]. In contrast, in cells transfected with the nsP3^HVD^ alone, the nsP3^HVD^ co-localized with both SG components eIF3b and G3BP1 in 93% of cells examined (Fig. 1a, c). These data indicate that HVD alone cannot suppress SG formation, but instead the domain associates with SG proteins.

To determine whether the arsenite-induced nsP3^HVD^-associated structures shared SG properties, we treated the cells with cycloheximide. Cycloheximide is an elongation inhibitor that traps mRNAs along with translation factors in polysomes, thus decreasing the availability of mRNAs for SG formation and resulting in SG disassembly (Fig. 1b; [9]). Therefore, *bona fide* SGs disappear upon cycloheximide treatment. After cycloheximide treatment, the nsP3^HVD^ no longer co-localized with translation factor eIF3b, suggesting that the nsP3^HVD^-associated structures formed upon arsenite treatment were SGs (Fig. 1b, c). On the other hand, the nsP3^HVD^ remained co-localized with G3BP1 in both unstressed and stressed conditions likely due to their ability to directly interact (Fig. 1, Fig. S1c). Taken together with work from others [36,40–42], these data suggest that expression of full-length nsP3 alters the association of individual SG components (e.g., release of eIF3b and retention of G3BP1 with nsP3), and it is likely that domain(s) other than HVD of nsP3 may be responsible for suppressing SG formation.

Given that the N-terminal MD possesses enzymatic activity to remove ADP-ribosylation [38,45] and the structural integrity of SGs is dependent on ADP-ribosylation [19], we hypothesized that the macrodomain of nsP3 (nsP3^MD^) could be responsible for suppressing SG formation. Similar to the full-length nsP3, overexpression of the MD fragment (1-162) inhibited the formation of SGs upon treatment with arsenite (Fig. 1a, c) or other SG inducers, such as mitochondrial stressor clotrimazole and endoplasmic reticulum stressor thapsigargin (Fig. S1d, e; [9]). However, unlike full-length nsP3, the nsP3^MD^ did not co-localize with G3BP1, presumably due to a lack of the G3BP-binding HVD (Fig. 1a). These data together suggest that while the nsP3^HVD^ is sufficient for formation of structures with SG components G3BP1 and eIF3b, nsP3^MD^ plays a critical role in regulating the association between these SG components.

### ADP-ribosylhydrolase activity of the nsP3 macrodomain suppresses SG formation

Next, we tested whether the ADP-ribose binding and hydrolase activities of nsP3^MD^ are responsible for suppressing SG formation (Fig. 2). Three classes of MD mutants were used [38]: (1) no ADP-ribose binding and little hydrolase activity (D10A, G32E, and G112E); (2) weak ADP-ribose binding and weak hydrolase activity (G32S); and (3) weak hydrolase activity but stronger ADP-ribose-binding than wild-type (WT) (Y114A). As with WT nsP3, all overexpressed full-length nsP3 MD mutants appeared in G3BP1-positive foci with or without arsenite (Fig. 2a, S2a). However, unlike WT, all full length nsP3 mutants co-localized with SG components G3BP1 and eIF3b upon arsenite treatment (Fig. 2a, b). Given that all tested mutants have deficiencies in ADP-ribosylhydrolase activity, their inability to prevent SG formation is likely due to the absence of this enzymatic activity.

**Figure 2.**
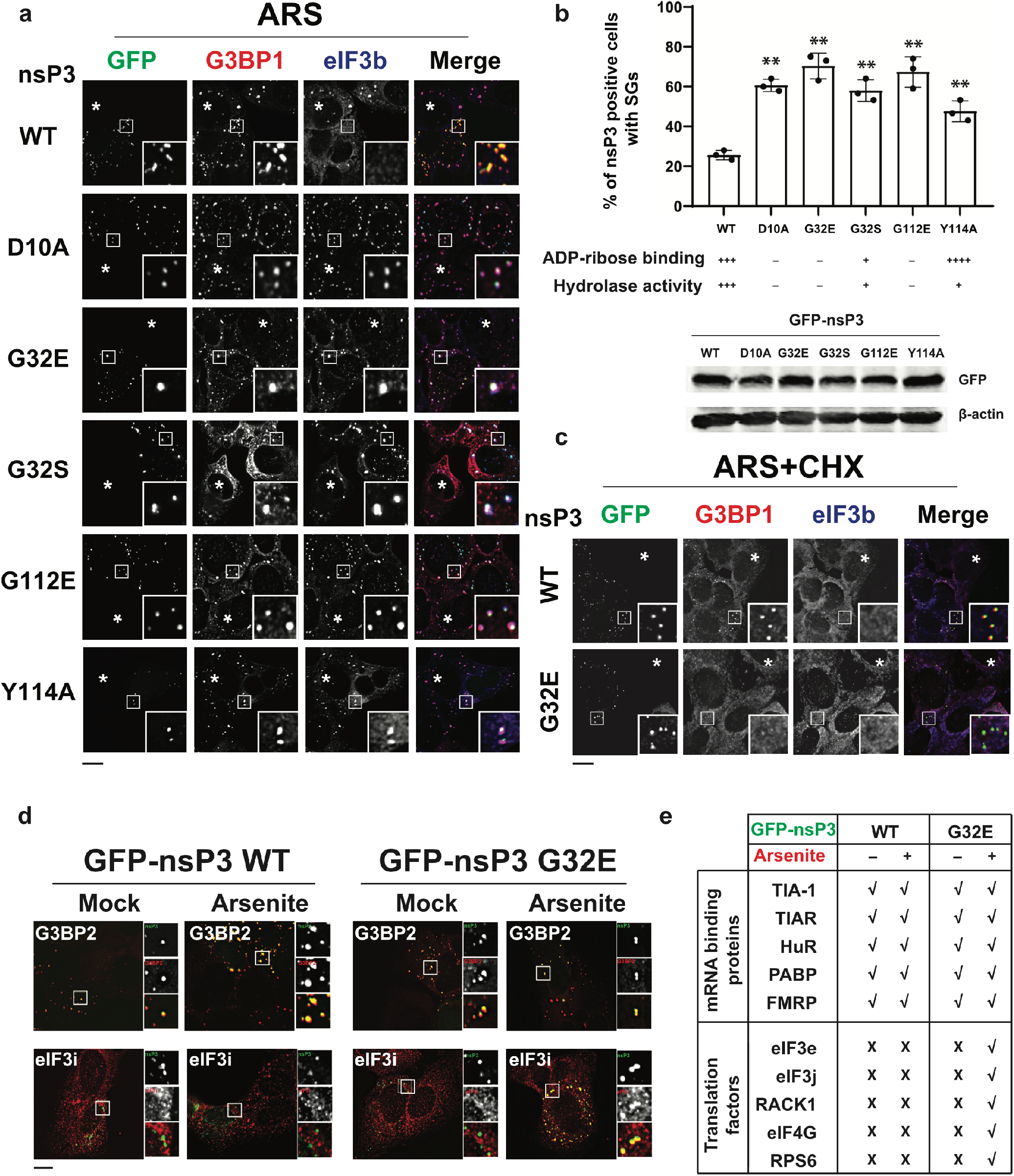
ADP-ribosylhydrolase activity of nsP3 suppresses SG formation. (a) U20S cells transfected with WT nsP3 or nsP3 with different point mutations (D10A, G32E, G32S, G112E, Y114A) were treated with 0.2 mM arsenite (ARS) for 30 min and immunostained for G3BP1 (red) and elF3b (blue). Asterisks indicate untransfected cells. (b) Bar graph indicates the percentage of nsP3-positive cells with SGs. Lower panels indicate ADP-ribose binding and hydrolase activity of the mutants based on McPherson et al., PNAS 2017, as well as their expression. **, p<0.01, two tailed, unpaired Student’s t test. (c) Cells were transfected with WT or G32E nsP3 and treated with 0.2 mM arsenite and 100 μg/ml cycloheximide (ARS+CHX) for 30 min. Cells were immunostained for G3BP1 (red) and elF3b (blue). Asterisks indicate untransfected cells. (d) U20S cells transfected with either GFP-tagged WT or G32E nsP3 (green) were mock-treated or treated with 0.2 mM arsenite for 30 min and immunostained for G3BP2 (red) and elF3i (red). (e) A table summarizes the immunofluorescence data in Supplementary Figure 3 on co-localization of nsP3 (WT or G32E) with different mRNA binding proteins or translation factors. Scale bar, 10 μm.

Using G32E as an example, we characterized further whether the foci formed by nsP3 mutants with G3BP1 and eIF3b share the properties of SGs. To test this possibility, cells expressing either the WT nsP3 or G32E mutant nsP3 were treated with arsenite and cycloheximide (*cf*. Fig. 1b). WT nsP3 remained co-localized with G3BP1, but not with eIF3b. On the contrary, while mutant nsP3 appeared as foci, eIF3b and the majority of G3BP1 did not (Fig. 2c). Therefore, mutant nsP3/G3BP1/eIF3b foci shared the cycloheximide-sensitive nature of canonical SGs. In addition, G32E expression also resulted in the co-localization of G3BP1 and eIF3b in response to other types of stress such as clotrimazole and thapsigargin (Fig. S2b, c), suggesting that the observed phenotype is common to multiple stressors that induce SGs.

SGs are condensates of stalled translation initiation complex along with proteins associated with the untranslated mRNAs. Given that the SG proteome is comprised of multiple protein-protein interaction networks [54,55], we tested whether ADP-ribosylhydrolase activity of nsP3 alters the co-localization of SG components in addition to G3BP1 and eIF3b in cells expressing either WT nsP3 (hydrolase-positive MD) or the G32E mutant (hydrolase-negative MD). The examined SG components included the RNA-binding proteins G3BP2, TIA1, TIAR, FMRP, HuR, and PABPC1, as well as the translation-related proteins eIF3e, eIF3i, eIF3j, eIF4G1 and RACK1, and RPS6 (Fig. 2d, e, and S3). As for G3BP1, WT nsP3 co-localized with all tested RNA-binding proteins in both untreated and arsenite-treated cells. On the other hand, as in the case of eIF3b, none of the translation-related proteins colocalized with WT nsP3 in either condition. However, the nsP3 mutant G32E co-localized with all tested SG components in arsenite-treated, but not untreated, cells. Taken together, these data suggest that the nsP3 ADP-ribosylhydrolyase activity regulate associations between SG components.

### SG formation is suppressed by ADP-ribosylhydrolase activity of nsP3 during viral infection

To determine whether the observed role of nsP3^MD^ ADP-ribosylhydrolase activity during arsenite-induced stress is relevant in a physiological context, we tested whether MD hydrolase mutation affects SG assembly in CHIKV-infected cells. Given that CHIKVs with nsP3 mutations (D10A, G32E, G112E) that severely compromise ADP-ribose binding and hydrolase activities are not viable [38], we focused on viable CHIKV mutants G32S and Y114A with diminished, but not absent, hydrolase activity. CHIKV WT and nsP3 mutants were examined 5.5 h after infection when nsP3 appears as foci in the absence of arsenite (Fig. S4a). To examine the ability of WT, G32S and Y114A nsP3s to suppress SG formation, infected cells were treated with arsenite for 0.5 h (a time point when ~95% of uninfected cells have developed SGs; data not shown). Consistent with experiments using nsP3 protein expression constructs (Fig. 2a, b), WT CHIKV infection suppressed SG formation, where only 27% of nsP3-positive cells had canonical (G3BP1/eIF3b double-positive) SGs (Fig. 3a, b).

**Figure 3:**
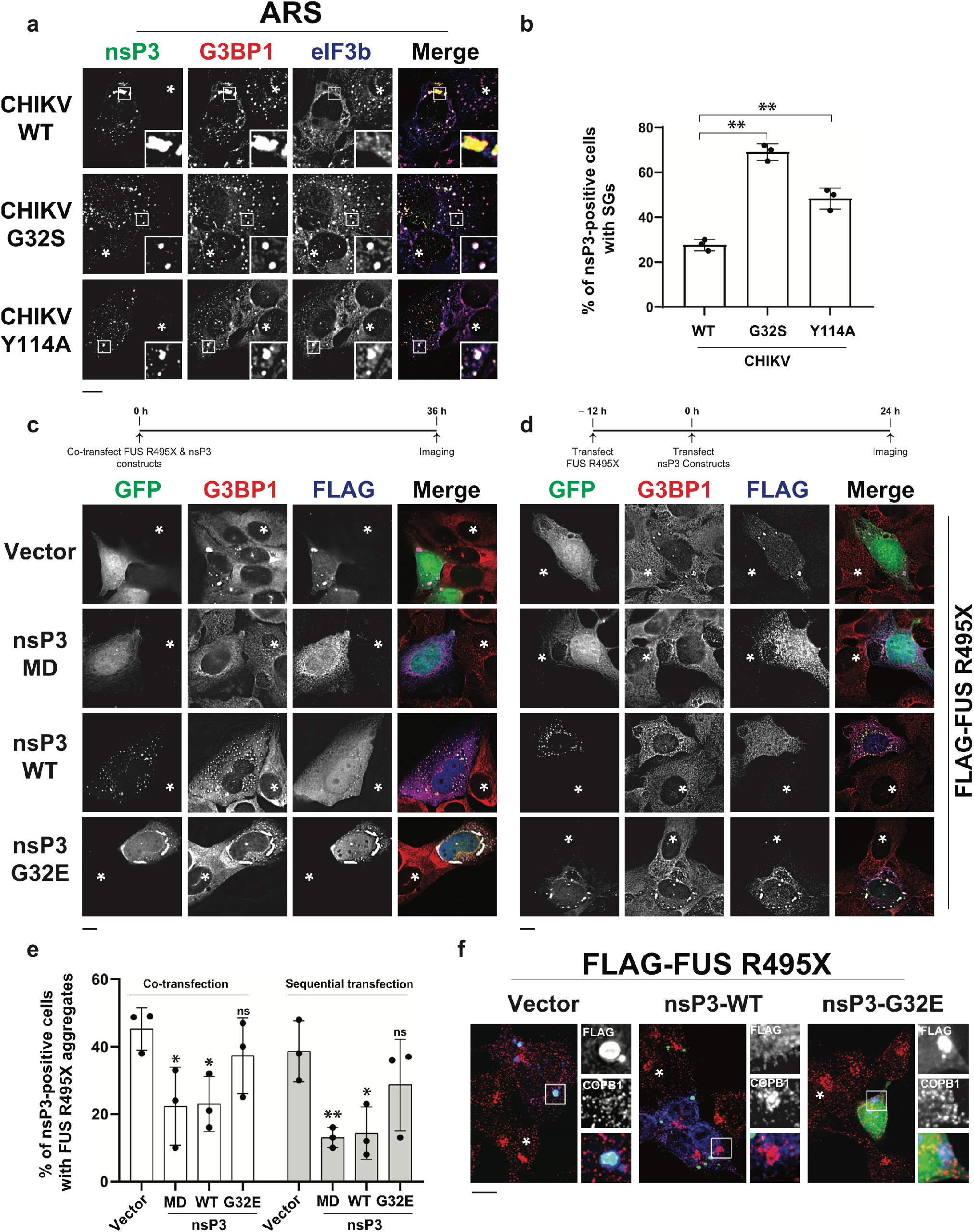
ADP-ribosylhydrolase activity of nsP3 suppresses formation of SGs during virus infection and SG-like aggregates upon expression of ALS-linked FUS R495X mutant. (a) U2OS cells were infected with CHIKV containing a WT, G32S or Y114A nsP3 at an MOI = 1. 5.5 hours post infection (hpi) cells were treated with 0.2 mM arsenite (ARS) for 30 min and immunostained for nsP3 (green), G3BP1 (red) and elF3b (blue). (b) Bar graph shows the percentage of virus-infected cells (nsP3-positive) with SGs. **, p<0.01, two tailed, unpaired Student’s t test. U2OS cells were either (c) co-transfected or (d) sequentially transfected (as indicated in the experimental scheme) with FLAG-tagged FUS R495X and either GFP vector, GFPtagged WT nsP3, G32E nsP3 or nsP3^MD^. At 36 h post-transfection, cells were processed for immunofluorescence, stained for FLAG (blue) and G3BP1 (red). (e) Bar graph shows the percentage of nsP3-positive cells with FUS R495X aggregates. Statistical comparison with vector control. *, p<0.05, **, p<0.01, two tailed, unpaired Student’s t test. (f) SH-SY5Y cells were transfected with FLAG-FUS R495X and either GFP vector, GFP-tagged WT nsP3 or G32E nsP3. Cells were processed as in (c) and stained for FLAG (blue) and COPB1 (red). Asterisks indicate (a) uninfected and (c-d, f) untransfected cells. Scale bar, 10 μm.

In contrast, 69% of G32S and 49% of Y114A nsP3-positive cells displayed canonical SGs (Fig. 3a, b). Similar colocalization of mutant nsP3, eIF3b and G3BP1 were observed with different SG inducers such as clotrimazole and thapsigargin (Fig. S4b-c), suggesting that the requirement of ADP-ribosylhydrolase activity for suppression of SG formation during viral infection is not stress-specific. To test whether these mutant nsP3/eIF3b/G3BP1 foci are sensitive to change in translation similar to SGs, we treated the infected cells with cycloheximide. Similar to the studies expressing nsP3 protein alone (Fig. 2c), the translation initiation factor eIF3b was diffusely distributed in the cytoplasm rather than in foci (Fig. S4d). However, mutant nsP3 and G3BP1 remained as foci during viral infection, suggesting that G3BP1 associates with additional viral proteins besides nsP3 in these foci (e.g., [56]). Taken together, these data suggest that the ADP-ribosylhydrolase activity of nsP3 can alter the association between individual SG components during viral infection.

### ADP-ribosylhydrolase activity of nsP3 suppresses the formation of and disassembles ALS mutant-mediated pathological SGs

Formation of aberrant SGs is a hallmark of several neurodegenerative diseases including amyotrophic lateral sclerosis (ALS) and frontotemporal dementia (FTD) [1,57]. Genetic mutations are commonly observed in RNA-binding proteins such as FUS *(e.g.* R495X), and overexpression of some ALS-related mutant RNA-binding proteins results in the formation of aggregates enriched in canonical SG components such as G3BP1 and translation initiation factors [58,59]. We hypothesized that the formation of FUS R495X-mediated SG-like aggregates might also be suppressed by nsP3 ADP-ribosylhydrolase activity. We co-transfected FLAG-tagged FUS R495X with either GFP alone, GFP-tagged nsP3 WT or GFP-nsP3 G32E into U2OS cells and examined the presence of FUS R495X-induced aggregates. Consistent with previous reports [48,49,60], overexpression of FUS-R495X itself formed SG-like aggregates, which were positive for G3BP1 and eIF3b, in ~50% of transfected cells (vector, Fig. 3c, e, data not shown). Intriguingly, co-transfection with full-length nsP3 WT, but not the point mutant G32E, decreased formation of these aggregates by two-fold (Fig. 3c, e). Moreover, the nsP3 G32E mutant co-localized with FUS R495X as well as SG markers G3BP1 and eIF3b (Fig. 3c and data not shown). Given that the full length nsP3 protein possesses the G3BP-binding HVD, the reduction of SG-like aggregates was accompanied by formation of foci containing both nsP3 and G3BP1 (Fig. 3c). On the contrary, co-transfection of nsP3^MD^ alone decreased formation of these aggregates by two-fold without the formation of nsP3/G3BP1 foci (Fig. 3c, e), further pointing to the importance of the MD ADP-ribosylhydrolase activity.

To determine whether the ADP-ribosylhydrolase activity can also cause the dissolution of pre-formed FUS R495X-induced aggregates, we transfected the FUS R495X construct 12 h prior to the nsP3 constructs (Fig. 3d). As in the case for co-transfection, transfection of WT nsP3 or MD alone, but not G32E nsP3, decreased the number of aggregates by at least 2.5-fold (Fig. 3d-e). These data suggest that the SG-like aggregates induced by FUS R495X can also be disrupted by nsP3 ADP-ribosylhydrolase activity.

Expression of FUS R495X in neuronal cells is reported to cause the dispersion of the protein COPBI—the coatomer beta subunit of the coat protein complex I involved in retrograde vesicular trafficking from Golgi and ER [61]. Intriguingly, expression of nsP3 WT, but not the G32E mutant, restored the dispersed pattern of COPBI in FUS R495X-transfected neuronal SH-SY5Y cells to the compact, juxtanuclear pattern observed in untransfected cells (Fig. 3f). As in U2OS, SG-like aggregates were induced by the expression of FUS R495X in these neuronal cells (Fig. S4e-f), and the number of aggregates were reduced upon co-transfection with nsP3 WT, but not G32E mutant (Fig. S4e-f). These data further suggest that the nsP3 ADP-ribosylhydrolase activity restores at least one cellular property disrupted by the FUS R495X mutant in addition to SG disassembly.

### nsP3 reduces ADP-ribosylation associated with the essential stress granule component G3BP1

Next, we examined the possible molecular targets for the ADP-ribosylhydrolase activity of nsP3 that altered SG composition and structural integrity. We focused on G3BP1 as a candidate because it is a direct binding partner of nsP3 [40,41,44,50–52], it is an essential component of SGs formed upon treatment with arsenite, clotrimazole and thapsigargin [10], it plays a critical role in virus infection [40,62], and it is required for the formation of SG-like aggregates associated with ALS [63–65]. Given that the binding of G3BP1 by the nsP3 HVD is proposed to be critical for SG disassembly [35,36] and that the ADP-ribosylhydrolase mutant of nsP3 has a decreased ability to suppress SG formation (Fig. 2a), we next tested whether the observed change in colocalization of SG components is due to the differential ability for nsP3 mutants to bind G3BP1 (Fig. 4a). GFP-tagged WT or mutants representative of three different classes (G32E, G32S, and Y114A) were transfected into cells, immunoprecipitated and probed for G3BP1. No differences were observed for the G3BP1 association with WT nsP3 and mutants tested, suggesting that ADP-ribosylhydrolase activity does not regulate the association between nsP3 and G3BP1.

**Figure 4:**
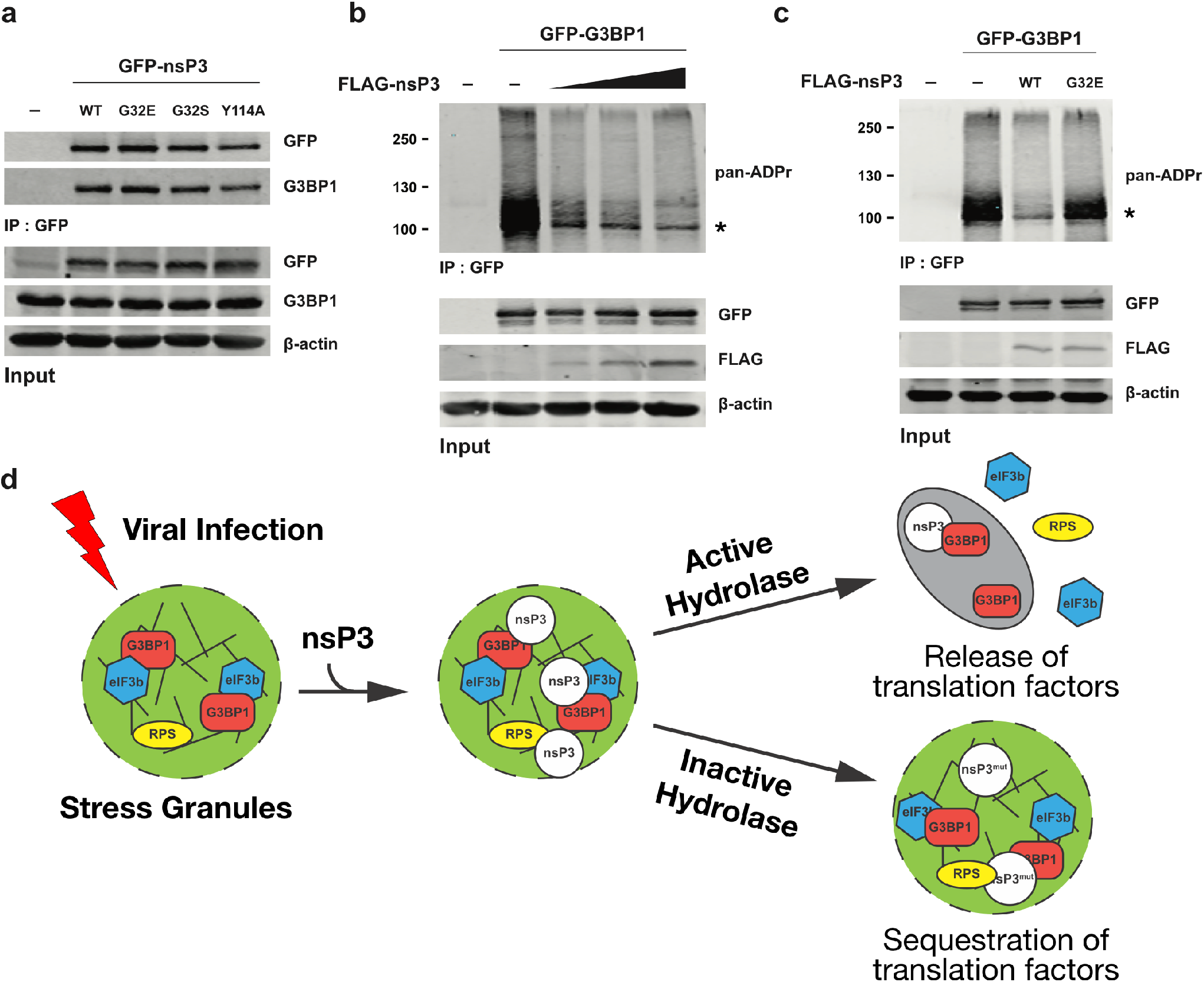
nsP3 reduces the ADP-ribosylation associated with the essential SG component G3BP1. (a) 293F cells were transfected with GFP vector or GFP-tagged WT nsP3, G32E nsP3, G32S nsP3, or Y114A nsP3. After 24 h transfection, cells were pelleted, lysed and immunoprecipitated using anti-GFP antibodies. The immunoprecipitates were blotted against against G3BP1 and GFP antibodies. (b) 293F cells were transfected with either GFP vector, GFP-tagged G3BP1 alone, or co-transfected with GFP-tagged G3BP1 with increasing concentration of FLAG-tagged nsP3^WT^. After 24 h transfection, cells were pelleted, lysed, immunoprecipitated using anti-GFP antibodies and immunoblotted with pan-ADPr reagent.(c) 293 cells were transfected with GFP vector, GFP-tagged G3BP1 alone or co-transfection of GFP-tagged G3BP1 with either FLAG-tagged WT or G32E nsP3. Cells were pelleted, lysed, immunoprecipitated using anti-GFP antibodies and immunoblotted with pan-ADPr reagent. Asterisks in panels b and c indicate the expected molecular weight of GFP-tagged G3BP1. (d) Model of nsP3-mediated SG disassembly. RPS, ribosomal proteins from 408 subunits.

Besides binding to nsP3, G3BP1 and its associated proteins are also ADP-ribosylated [19] and, therefore, might be targets of nsP3 ADP-ribosylhydrolase activity. To test this possibility, we expressed GFP-G3BP1 alone or co-expressed it with increasing amounts of FLAG-tagged nsP3. GFP-G3BP1 was then immunoprecipitated and probed for ADP-ribosylation. With GFP-G3BP1 expression alone, a prominent signal of ADP-ribosylation appeared at and above the expected molecular weight of GFP-G3BP1. The signal appeared as a smear because of the heterogeneous number of ADP-ribose units added onto G3BP1 and its associated proteins. Upon co-expression of WT full-length nsP3, the ADP-ribosylation signal associated with G3BP1 was reduced in a dose-dependent manner (Fig. 4b). Such reduction of signals was not observed when the nsP3 G32E mutant was co-expressed (Fig. 4c), suggesting that the ADP-ribosylhydrolase activity of nsP3 is responsible for reducing ADP-ribosylation associated with the essential SG component G3BP1.

## DISCUSSION

The presence of microscopically visible SGs is a manifestation of the underlying dynamic equilibrium of factors that control its rates of assembly and disassembly at a sub-microscopic level. Emergent data indicate that this equilibrium can be altered through post-translational modifications, whereby enzymes with opposing activities that add and remove these modifications regulate the dynamics of SG integrity [20]. Here, we report that the SG integrity is regulated by the MD of alphaviral protein nsP3 through its ability to enzymatically remove ADP-ribosylation. Expression of the wild-type nsP3 MD alone is sufficient to suppress SG formation. The requirement for ADP-ribosylhydrolase activity was demonstrated by the loss of ability to suppress SG formation by five different mutants with impaired enzyme activity. Though it is possible that activities other than ADP-ribosylhydrolase possessed by the nsP3 MD may be responsible for suppressing SG formation, the fact that mutations at different sites in the MD result in the same phenotype make such a scenario less likely. Importantly, the expression of WT nsP3, but not the ADP-ribosylhydrolase-deficient mutant nsP3s, suppresses the formation of SGs in response to multiple types of stress including oxidative stress (arsenite), mitochondrial stress (clotrimazole) and ER stress (thapsigargin). Suppression was observed both by ectopic expression of nsP3 through transfection and natively in the context of CHIKV infection. Given that the MD and its ADP-ribosylhydrolase activity are conserved across all macrodomain-containing RNA viruses [66,67], including coronaviruses, it is possible that these viruses suppress SG formation through a common mechanism by removing ADP-ribosylation.

For alphaviruses, SGs are first induced in the early stage of infection and subsequently disassembled [32,34]. It was previously proposed that the disassembly is mediated through the sequestration of the essential SG component G3BP1/2 by associating with the C-terminal HVD domain of nsP3 [35,36,41]. Though we confirmed the association between nsP3 and G3BP1, our data show that the expression of the HVD domain alone does not suppress SG formation. It is therefore unlikely that nsP3 causes SG disassembly through HVD sequestration of G3BP1/2 alone. Yet, G3BP1/2 and its association with the nsP3 HVD are required for CHIKV replication [40,62]. We note, however, that the HVD is composed of low-complexity regions [39], which are increasingly appreciated for their role in mediating the assembly of non-membranous structures [6–8]. Here, we instead propose a model of how nsP3 mediates SG disassembly, wherein the C-terminal HVD promotes the association between nsP3 and SG proteins so that the N-terminal MD can act locally to remove ADP-ribosylation in SGs. As an increasing amount of nsP3 is expressed over the course of infection, a higher level of ADP-ribosylhydrolase activity will suppress the rate of SG assembly and thereby shift the dynamic equilibrium towards the disappearance of SG in the late stage of the replication cycle.

One may wonder: what could be the possible roles for nsP3 ADP-ribosylhydrolase to disassemble SGs—i.e., condensates of stalled translation initiation complexes—during the late stage of alphavirus infection? The answer may be revealed by examining what happens to cells upon the expression of ADP-ribosylhydrolase-deficient mutants. The expression of these mutants leads to the formation of SGs that appear non-canonical because of the nsP3 presence. Yet, these non-membranous structures share the same cycloheximide sensitivity as *bona fide* SGs and contain at least 14 canonical SG components. Intriguingly, expression of enzymatically active nsP3 results in the segregation of all tested SG-associated RNA-binding proteins (G3BP1, G3BP2, TIA-1, TIAR, HuR, PABP and FMRP) from all tested 7 SG-associated translation factors (eIF3b, eIF3i, eIF3e, eIF3j, RACK1, eIF4G1 and RPS6). In contrast, all of these proteins are aggregated upon stress in the absence of ADP-ribosylhydrolase activity, suggesting that ADP-ribosylation is involved in determining the composition of SGs. Recent data indicate that PAR can recruit proteins to SGs through PAR-binding motifs [25,26,29]. It would be of interest to explore whether PAR plays a role in re-assortment of proteins in non-membranous structures in general. Our analyses further revealed that the essential SG component G3BP1 is one of the nsP3 ADP-ribosylhydrolase targets. However, it is unclear whether the observed re-assortment of SG composition is mediated by ADP-ribosylation of G3BP1 and/or other targets within SGs, which warrants further investigation. The observed pattern of specific protein retainment suggests that some protein-protein interaction networks within SGs are mediated by PAR, with the possibility that the ADP-ribosylhydrolase activity of nsP3 actively promotes the release from SGs of translation factors needed for viral protein translation (Fig. 4d). Such a possibility is consistent with recent data showing that nsP3 ADP-ribosylhydrolase activity is critical for switching from host translation to translation of the viral structural proteins during the later stages of viral replication [68,69]. This re-distribution of host translation initiation factors and ribosomal proteins resulting from the shutoff of host translation may be required for virus production at later stages of infection [32,34].

Informatics analyses revealed that RNA granule proteins are enriched for low complexity regions and these proteins are preferentially modified by ADP-ribosylation and that PAR-binding proteins are also enriched for low-complexity regions [27,70–72]. Compared with other protein modifications that regulate SG dynamics, ADP-ribosylation is unique in that the polymeric form (i.e., PAR) is able to seed the low-complexity region-containing proteins to form non-membranous structures *in vitro* [25–28,70]. Our data suggest that the nsP3 ADP-ribosylhydrolase likely degrades the PAR that seeds SGs in cells, with G3BP1 as a possible ADP-ribosylated substrate (cf. Fig. 4b, c). The complete removal of PAR from ADP-ribosylated proteins requires two steps: the degradation of the polymeric chain down to single ADP-ribose units conjugated to proteins, followed by the hydrolysis of the final, proximal ADP-ribose groups from proteins [73–75]. The alphaviral nsP3 MD alone efficiently hydrolyzes single ADP-ribose groups from ADP-ribosylated proteins *in vitro* but inefficiently removes the PAR chains [38,45]. However, it is possible that the nsP3 MD may partner with other protein(s) to process the PAR chains in cells. For example, the hepatitis E viral MD hydrolyzes PARylated proteins efficiently *in vitro* through association with another viral protein that possesses helicase activities [47]. Notably, several host helicases associate with alphaviral nsP3 in infected cells [39], and future experiments will help address whether or not any of them are involved in partnering with nsP3 to remove PAR. Alternatively, our previous studies identified PARG—an endogenous enzyme that cleaves the ribose– ribose bonds between ADP-ribose units—as localized in SGs, where its expression level controls the assembly and disassembly of SGs [19]. Therefore, another model is that PARG mediates the breakdown of PAR in SGs, followed by nsP3 ADP-ribosylhydrolase providing the critical step to completely remove the final ADP-ribose from modified proteins.

SGs and non-membranous condensates are observed in many physiological and pathological contexts in virus infection, cancers and neurodegenerative diseases [1,12–14]. Preclinical studies showed that PARP inhibitors reduce the number of non-membranous pathological aggregates in models of neurodegenerative disease [26,30,76] and PARG knockout flies revealed progressive neurodegeneration with observable cytoplasmic aggregates [77]. Consistent with these findings, our data showed that expression of wild-type nsP3, but not an ADP-ribosylhydrolase-deficient nsP3 mutant, can reduce the number of SG-like aggregates formed upon expression of the FUS protein mutant R495X, which is linked to an aggressive form of ALS. These data altogether suggest possible therapeutic options for reduction of non-membranous protein aggregates by lowering PAR levels, either through inhibiting PARP or increasing the ADP-ribosylhydrolase activity.

## METHODS

### Cell culture, chemicals and transfection

U2OS and HeLa cells were obtained from ATCC and 293F cells from Invitrogen. Cells were maintained in DMEM medium (Gibco) containing 10% heat-inactivated FBS (Gibco, Life Technologies). In all experiments, plasmids were transfected using JetPrime from Polyplus (U2OS) or 293fectin from Gibco (293F) as per manufacturer’s protocols. The following drugs were used for stress induction: Arsenite (Sigma, S7400); Clotrimazole (Sigma, C6091); Thapsigargin (Sigma, T9033); Cycloheximide (Sigma, C7698).

### Viruses

CHIKV 181/25 WT and mutant strains were prepared as described previously [69]. Viral stocks were grown in BHK21 cells and assayed by plaque formation in Vero cells.

### Immunoblot analysis

Cells were lysed in RIPA buffer (50 mM Tris-Cl pH 8.0, 150 mM NaCl, 0.1% SDS, 1% NP-40, 1 mM EDTA, containing proteinase inhibitors 5 mM NaF, 1 mM PMSF) for 15 min on ice followed by centrifugation at high speed for 15 min, 4°C. Protein samples were acetone-precipitated for at least 1 h at −20°C. Precipitates were centrifuged at 13,000 rpm, 4°C for 15 min, and the air-dried pellets were then diluted in 1X SDS sample buffer. The samples were resolved in SDS-PAGE gel and blotted with appropriate primary antibodies (Table S2).

### Immunofluorescence

~4×10^4^ (U2OS) or ~1×10^5^ (HeLa) cells grown on coverslips were treated with the indicated stressors. Following the stress treatment, cells were washed twice with 1X PBS, fixed with 4% paraformaldehyde for 15 min, permeabilized with ice-cold methanol for 10 min and washed twice with 1X PBS. The cells were then blocked with 5% normal horse serum in 1X PBS containing 0.02% sodium azide (NHS/PBSA) for 1 h at RT. All primary antibodies (Table S2) were diluted in blocking buffer and incubated with cells overnight at 4°C, followed by three washes with 1X PBS, 10 min each. Next, appropriate secondary antibodies were diluted in blocking buffer along with Hoechst (1 μg/ml), incubated with the cells for 1 h at RT, washed three times with 1X PBS and the coverslips were mounted on glass slides using Prolong Gold. All experiments were performed at least thrice.

### Image quantitation

For all quantitation, random 40X fields were chosen and ~120 cells were counted per condition. To quantify the number of SGs in cells transfected with various GFP-tagged nsP3 constructs, cells with eIF3b foci in GFP-positive cells were considered as SG-positive. To quantify the number of SGs in virus infection, eIF3b foci in nsP3-stained cells were considered as SG-positive. For the number of FUS495X aggregates in U2OS and SH-SY5Y cells, cells expressing both FLAG (FUS495X) and GFP (vector/nsP3) were counted for the presence of microscopically visible aggregates. All experiments were repeated three independent times.

### Immunoprecipitation

293F cells were spun down at 400 xg for 3 min at 4°C, washed once with cold 1X PBS, pelleted and lysed in cold lysis buffer (CLB) (50 mM HEPES pH 7.4, 150 mM NaCl, 1 mM MgCl_2_, 1 mM EGTA, 1% TRITON X-100 supplemented with 1 mM NaF, 1 mM PMSF, 1 mM ADP-HPD). Cell lysates were mixed at 4°C for 15 min, spun down for 15 min at 13,000 rpm and the supernatant fluid was collected in a new tube. The cleared lysates were added to pre-incubated anti-GFP (3E6, Invitrogen)– DYNA magnetic beads (10004D, Invitrogen) complex and incubated for 2 h at RT. The beads were washed once with CLB, twice with high-salt CLB (300 mM NaCl), followed by a final wash with CLB. The precipitates were then boiled with 1X SDS sample buffer for 10 min at ~85°C. All experiments were performed at least thrice.

### Statistical analysis

Data are presented as mean ± SD and groups compared using two-tailed unpaired Student’s t test. *p* < 0.05 was considered statistically significant. All statistical analyses were performed using Graphpad Prism8.

## ACKNOWLEDGMENTS

We thank Drs. Phillip Sharp, Nancy Kedersha and Lucas Reineke and members of the Leung lab for their critiques of the manuscript and Dr. Rachy Abraham for supplying the viruses used. We thank Dr. Ted Dawson for SH-SY5Y cells and Dr. Steve McKnight for FLAG-FUS R495X construct. This work was supported by a Johns Hopkins Catalyst Award (A.K.L.L), pilot grants from the Johns Hopkins University School of Medicine Sherrilyn and Ken Fisher Center for Environmental Infectious Disease (A.K.J., D.E.G., A.K.L.L.) and in part by R01GM104135 (A.K.L.L.).

## AUTHOR CONTRIBUTIONS

A.K.J and A.K.L.L. conceived the project; A.K.J., D.E.G. and A.K.L.L. designed the experiments; A.K.J conducted the experiments; A.K.J. and A.K.L.L. wrote the paper with input from D.E.G.

---

**Table S1.** Number of copies of G3BP1 and G3BP2 per cell.

|  | U2OS<br>Beck M *et al.*<br>Mol Syst Biol 2011 | HeLa<br>Nagaraj *et al.*<br>Mol Syst Biol 2011 | HeLa<br>Hein *et al.*<br>Cell 2015 | HepG2<br>Wiśniewski *et al.*<br>J Proteomics 2016 |
| --- | --- | --- | --- | --- |
| **G3BP1** | 431,000 | 1,300,000 | 751,796 | 432,865 |
| **G3BP2** | 52,200 | 390,000 | 990,184 | 182,332 |
| Median protein abundance | 17,718 | 42,057 | 28,935 | 11,703 |

**Table S2.** Antibodies used in this study

| Antibody | Catalog | Dilution<br>Immunoblot | Dilution<br>IF | Source |
| --- | --- | --- | --- | --- |
| G3BP1 | sc-365338 | 1:500 | 1:500 | Santa Cruz |
| G3BP2 | A302-040A-M | 1:1000 | 1:500 | Bethyl |
| eIF3b | SC-16377 | 1:1000 | 1:200 | Santa Cruz |
| nsP3 |  | 1:10000 | 1:5000 | Andres Merits |
| eIF3i | sc-374156 |  | 1:500 | Santa Cruz |
| eIF3e | A302-985A |  | 1:500 | Bethyl |
| eIF3j | 10439-1-AP |  | 1:500 | ProteinTech |
| RACK1 | sc-17754 |  | 1:200 | Santa Cruz |
| eIF4G1 | SC-11373 |  | 1:400 | Santa Cruz |
| RPS6 |  |  | 1:250 | Santa Cruz |
| TIA1 | SC-1751 |  | 1:250 | Santa Cruz |
| TIAR | sc-1749 |  | 1:250 | Santa Cruz |
| FMRP | ab27455 |  | 1:500 | Abcam |
| HuR | SC-5261 |  | 1:250 | Santa Cruz |
| PABP | ab21060 |  | 1:400 | Abcam |
| FLAG | F1804 | 1:1000 | 1:1000 | Sigma |
| FLAG | F7425 | 1:1000 | 1:1000 | Sigma |
| GFP | 11814460001 | 1:500 |  | Roche |
| actin | A1978 | 1:1000 |  | Sigma |

**Supplementary Figure 1:**
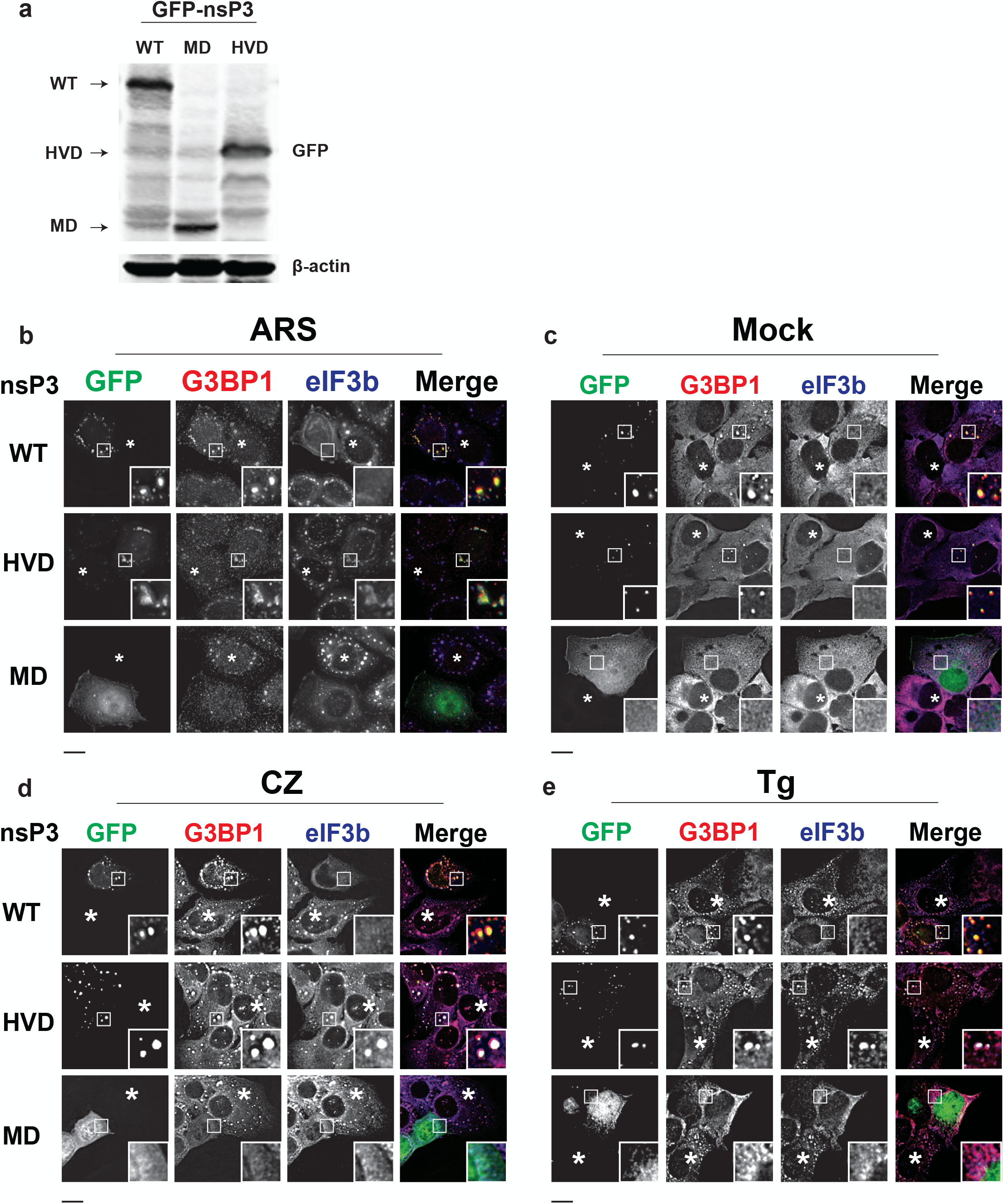
nsP3 macrodomain suppresses SG formation under different conditions, related to Figure 1. (a) Expression level of nsP3^WT^, nsP3^HVD^ and nsP3^MD^ in Figure 1a. (b) HeLa cells were transfected with GFP-tagged WT nsP3, nsP3^HVD^ or nsP3^MD^ and treated with 0.2 mM arsenite (ARS) for 30 min. Cells were then fixed and immunostained for SG markers G3BP1 (red) and eIF3b (blue). Though the majority of nsP3^MD^-expressing HeLa cells still contain some SGs as defined by the colocalization of G3BP1 and eIF3b, the size of these foci is significantly smaller compared with untransfected cells. (c-e) U2OS cells transfected with GFP-tagged WT nsP3, nsP3^HVD^ or nsP3^MD^ were either (c) mock-treated or treated with (d) 40 μM clotrimazole (CZ) or (e) 2 μM thapsigargin (Tg) for 30 min and immunostained for G3BP1 (red) and eIF3b (blue) antibodies. Asterisks indicate untransfected cells. Scale bar, 10 μm.

**Supplementary Figure 2:**
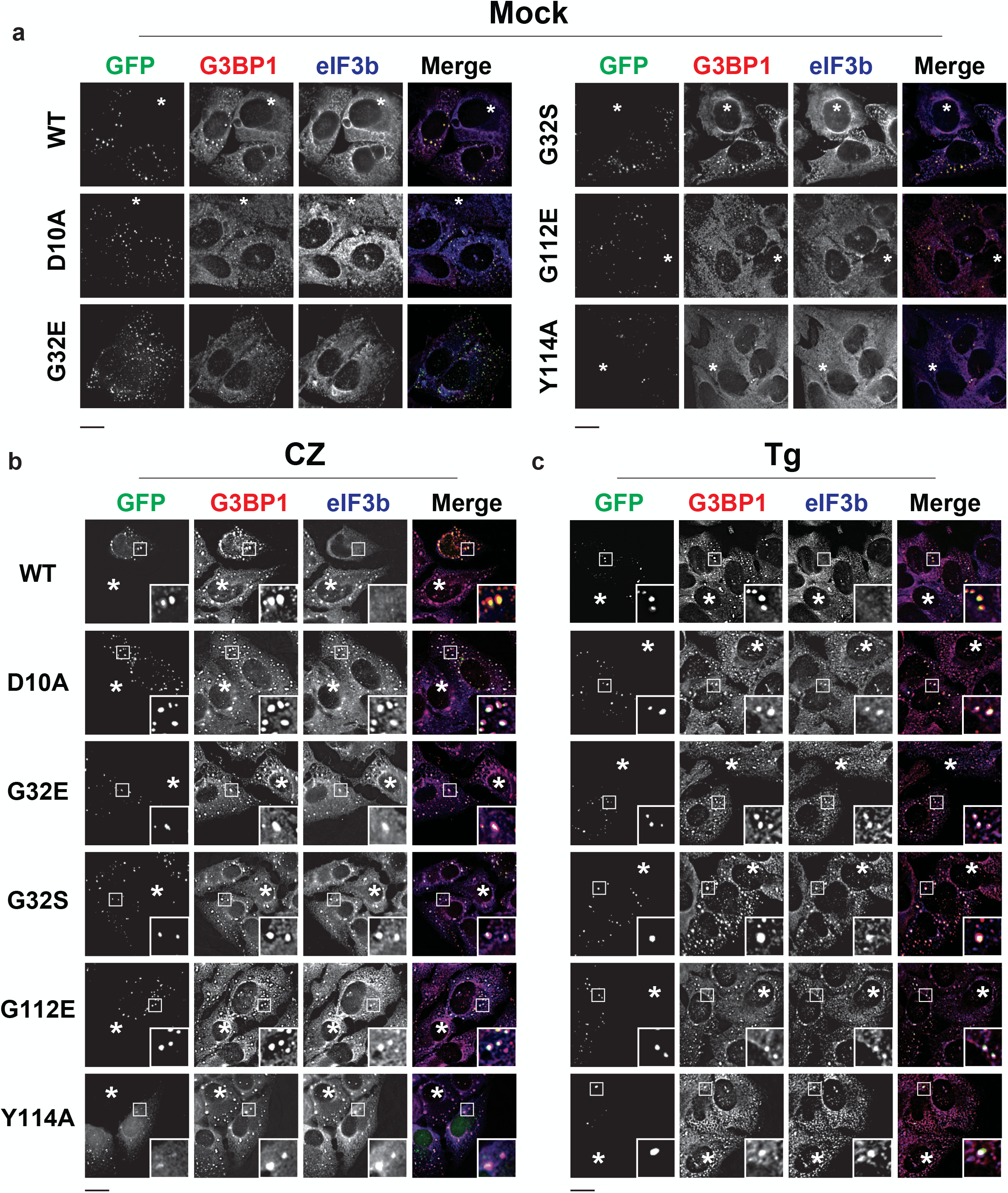
ADP-ribosylhydrolase activity of nsP3 suppresses SG formation under different conditions, related to Figure 2. U2OS cells transfected with WT nsP3 or point mutants D10A, G32E, G32S, G112E or Y114A were either (a) mock-treated or treated with (b) 40 μM clotrimazole (CZ) or (c) 2 μM thapsigargin (Tg) for 30 min or and stained for G3BP1 (red) and eIF3b (blue) antibodies. Asterisks indicate untransfected cells. Scale bar, 10 μm.

**Supplementary Figure 3:**
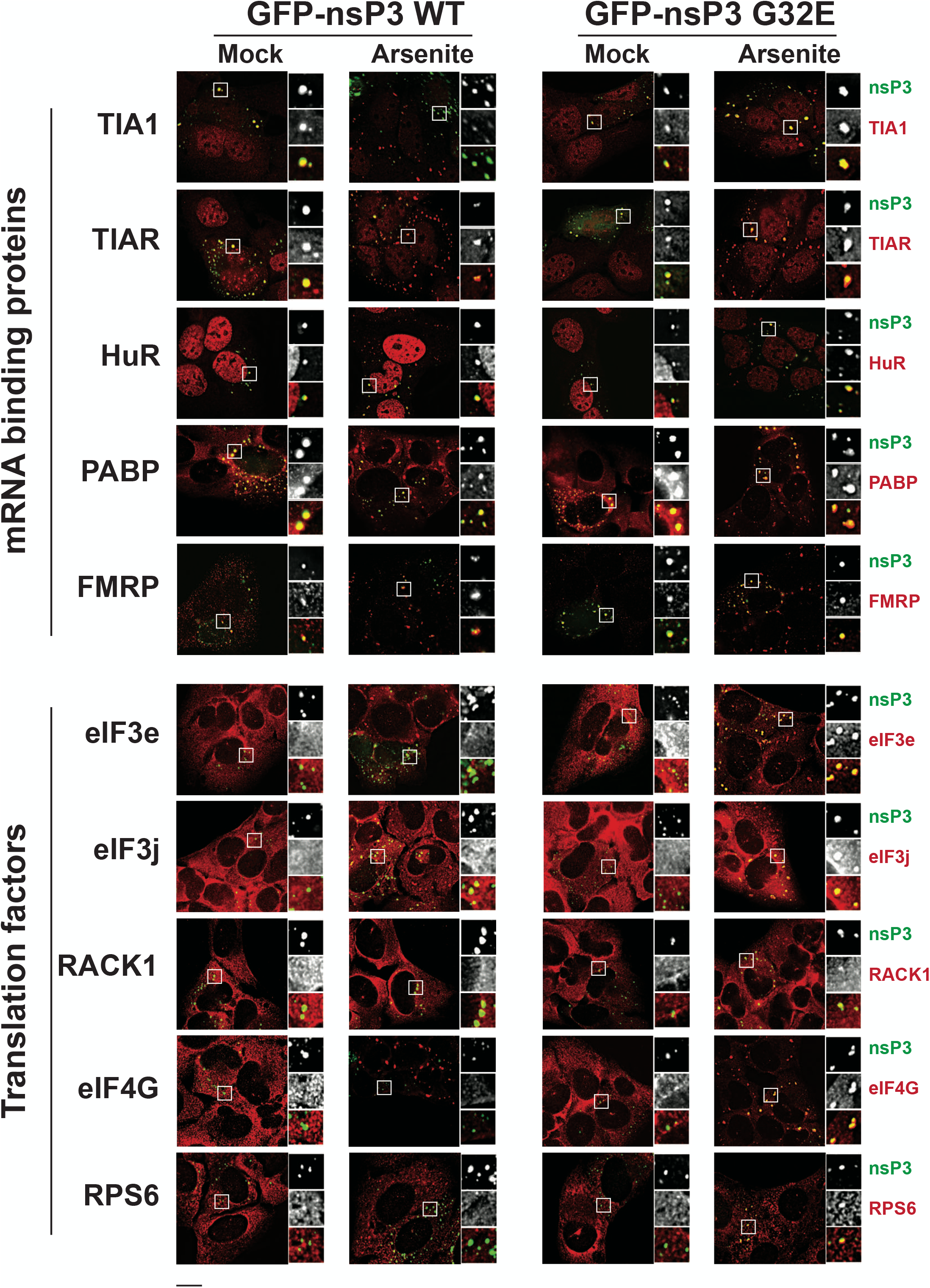
nsP3 differentially associates with SG components, related to Figure 2. (a) U2OS cells transfected with WT or G32E nsP3 were mock-treated or treated with 0.2 mM arsenite for 30 min and immunostained for the indicated proteins. Scale bar, 10 μm.

**Supplementary Figure 4:**
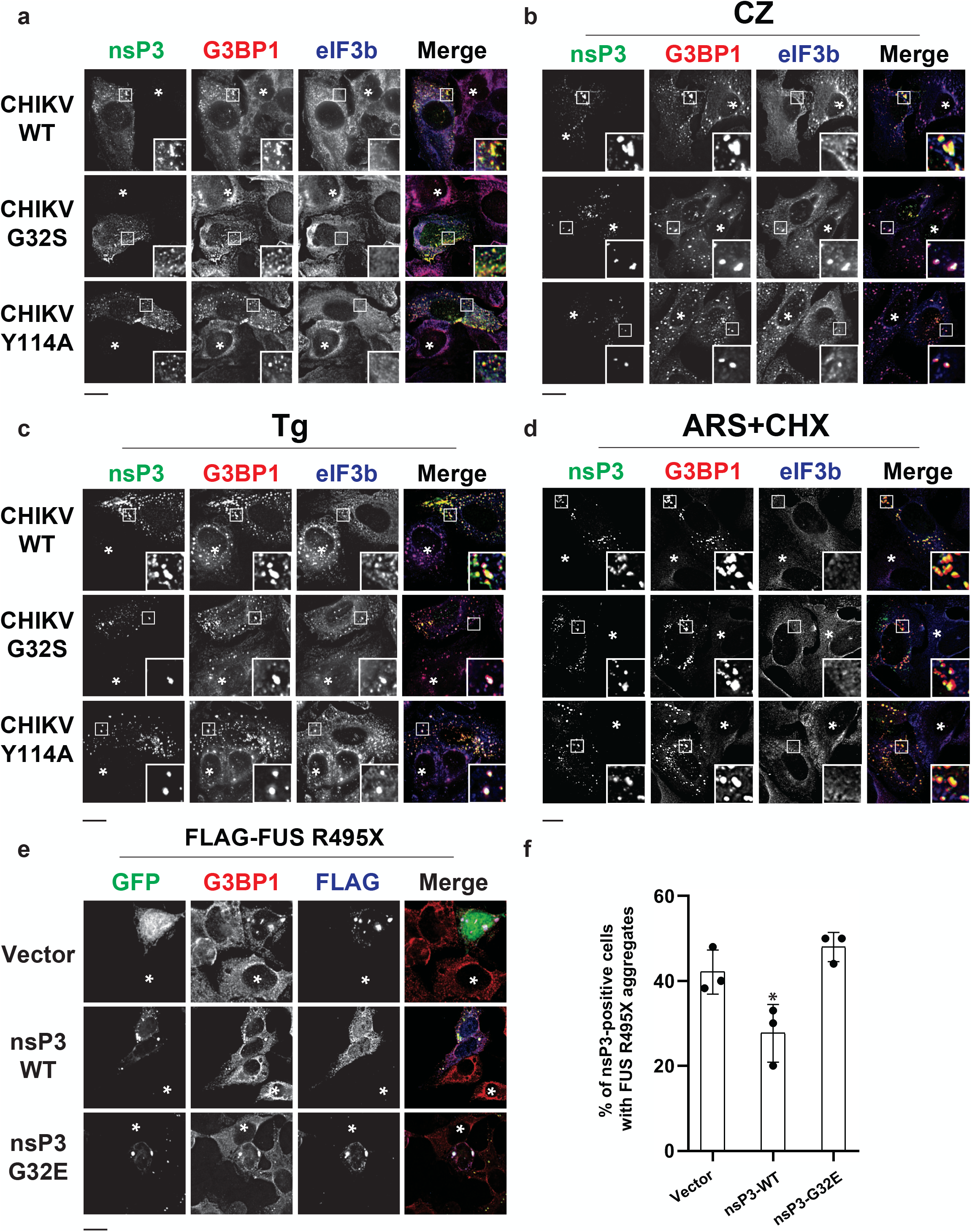
ADP-ribosylhydrolase activity of nsP3 suppresses SG formation during viral infection under different conditions and SG-like aggregates upon expression of ALS-linked FUS R495X mutant in neuronal SH-SY5Y cells, related to Figure 3. U2OS cells were infected with CHIKV expressing WT, G32S or 114A nsP3 at MOI = 1. (a) 6 hpi cells were fixed or (b-d) at 5.5 hpi, cells were treated with (b) 40 μM clotrimazole (CZ) or (c) 2 μM thapsigargin (Tg) (d) co-treated with 0.2 mM arsenite and 100 μg/ml cycloheximide (ARS+CHX) for 30 min and immunostained for nsP3 (green), G3BP1 (red) and eIF3b (blue). (e) SH-SY5Y cells were transfected with FLAG-FUS R495X and either GFP vector, GFP-tagged WT or G32E nsP3. At 36 h post-transfection, cells were processed for immunofluorescence, stained for FLAG (blue) and G3BP1 (red). *, p<0.05, two tailed, unpaired Student’s t test. Asterisks indicate (a-d) uninfected and (e) untransfected cells. (f) Bar graph shows the percentage of cells with aggregates in FLAG-FUS R495X transfected cells. Scale bar, 10 μm.

